# HUMAN-LIKE SIALOME REMODELING LINKS HYPERGLYCEMIA TO COLORECTAL CANCER PROGRESSION

**DOI:** 10.64898/2026.09.04.739806

**Authors:** Ronan C. M. Santos, Giulia S. Ferreira, Ana Luiza Lopes, Vanessa H. N. da Silva, Agata C. Fonseca, Miguel Fontes, César S. Bastos, Christina Maeda Takiya, João C. Machado, Miriam B. F. Werneck, Wagner B. Dias, Frederico Alisson-Silva, Adriane R. Todeschini

## Abstract

The evolutionary loss of cytidine monophosphate–N-acetylneuraminic acid hydroxylase (CMAH) abolished endogenous N-glycolylneuraminic acid (Neu5Gc) synthesis in humans, reshaping the sialic acid at the cell surface glycans and influencing immune and metabolic processes. However, whether a human-like sialome modulates the impact of metabolic stress on colorectal cancer (CRC) progression remains unclear.

Here, we investigated the diabetes–sialome–tumor axis using a spontaneous CRC model combining conditional *Apc* mutation with *Cmah* deficiency, thereby recapitulating the human sialic acid repertoire. Under euglycemic conditions, CPC-*Apc Cmah^-/-^* mice exhibited reduced polyp numbers and tumor burden compared with WT mice, accompanied by remodeling of the tumor immune microenvironment, including increased tumor-infiltrating leukocytes, enrichment of B cells, and reduced PD-1 expression in B cells and cytotoxic CD8^+^ T cells.

Induction of chronic hyperglycemia with low-dose streptozotocin revealed a striking genotype-specific effect. Despite comparable hyperglycemia between genotypes, tumor progression was dramatically exacerbated exclusively in *Cmah^-/-^* mice, with increased tumor burden, accelerated lesion development, and progression toward high-grade dysplasia. Histopathological analyses further revealed increased mesenchymal expansion and close resemblance to colorectal tumors from diabetic patients.

Together, these findings demonstrate that a human-like sialome increases susceptibility to metabolic stress and identify CPC-*Apc Cmah^-/-^* mice as a translationally relevant model for investigating diabetes-associated colorectal carcinogenesis.

**Graphical abstract:** 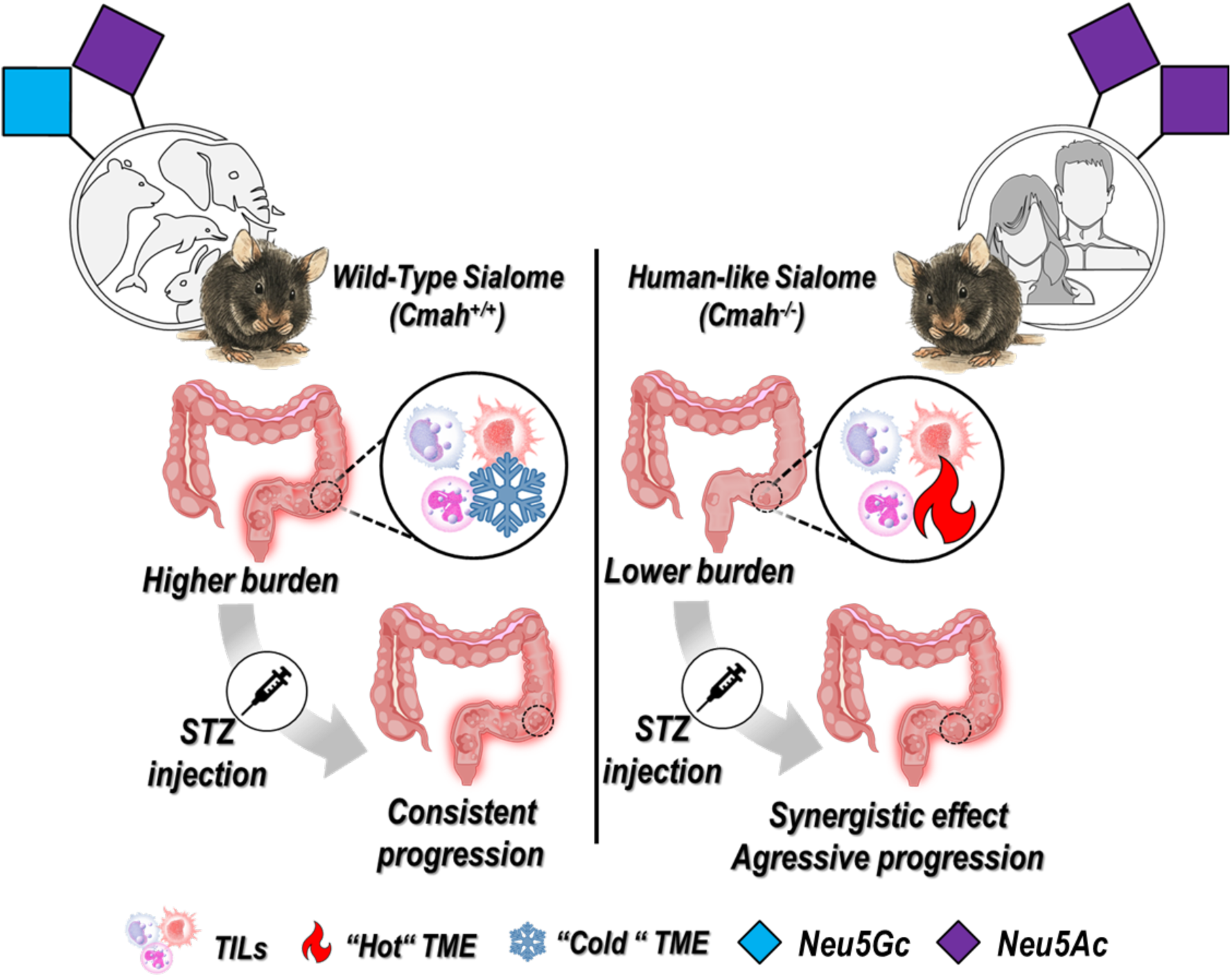

**Highlights:**

- CMAH loss reshapes the sialome and suppresses spontaneous colorectal tumorigenesis;
- Neu5Ac-dominant glycocalyx modifies tumor-infiltrating immune cell populations;
- Hyperglycemia selectively exacerbates tumor progression in *Cmah*^-/-^ mice;
- Diabetic *Cmah*^-/-^ tumors shift toward high-grade dysplasia with mesenchymal expansion;
- *Cmah*^-/-^ CPC-*Apc* model faithfully recapitulates diabetes-associated CRC progression in humans;
- Loss of CMAH increases susceptibility to hyperglycemia-driven colorectal cancer progression.

## Introduction

Colorectal cancer (CRC) remains one of the leading causes of cancer-related mortality worldwide [1–3], and its incidence and lethality are further exacerbated by metabolic comorbidities, particularly diabetes mellitus and chronic hyperglycemia [4–6]. Large clinical cohorts consistently report that diabetic patients exhibit higher CRC incidence, faster disease progression, increased recurrence, and reduced survival compared with non-diabetic individuals [5, 7]. Although these epidemiological associations are well established, the molecular mechanisms by which systemic dysglycemia translates into local tumor advantage remain incompletely understood.

Hyperglycemia profoundly influences tumor biology by promoting anabolic metabolism, reshaping immune responses, and reprogramming the tumor microenvironment [8–10]. Elevated glucose availability enhances CRC aggressiveness and promotes immune evasion [10–12] potentially through aberrant glycosylation, since glycan biosynthesis is highly sensitive to metabolic flux through the hexosamine biosynthetic pathway (HBP). Increased HBP activity elevates UDP-GlcNAc availability, leading to altered glycosylation patterns, including enhanced sialylation [9–11, 13, 14]. In CRC, increased sialylation has been associated with malignant progression, immune escape, and poor prognosis [15, 16].

Among glycan modifications, terminal sialic acids occupy a central position at the tumor–host interface. Sialic acids regulate immune checkpoints, receptor clustering, and interactions with sialic acid–binding immunoglobulin-like lectins (Siglecs), thereby shaping immune surveillance and inflammatory responses [15]. Due to an evolutionary loss of the Cytidine monophosphate-N-acetylneuraminic acid (CMP-Neu5Ac) hydroxylase (CMAH) enzyme, humans are unable to synthesize N-glycolylneuraminic acid (Neu5Gc) and instead exclusively express N-acetylneuraminic acid (Neu5Ac) [17, 18]. In contrast, most mammals, including *Mus musculus* mice, retain functional CMAH and endogenously produce Neu5Gc. This fundamental divergence creates a critical translational gap in modeling human glyco-immune interactions in cancer and metabolic disease.

Accumulating evidence indicates that CMAH loss has far-reaching consequences for immune regulation, inflammation, and metabolic homeostasis. CMAH-deficient (*Cmah*^-/-^) mice display enhanced immune reactivity, altered macrophage function, and increased inflammatory tone [19, 20], as well as higher susceptibility to glucose intolerance and metabolic dysregulation [17, 21]. In parallel, recent studies have linked aberrant Neu5Gc biology, including increased transcription of the *CMAH* pseudogene and dietary Neu5Gc incorporation, to increased aggressiveness of gastrointestinal tumors [22, 23]. Together, these findings suggest that the human-specific sialome fundamentally reshapes the tumor interface, both metabolically and immunologycally.

Based on our latest work, we hypothesized that hyperglycemia promotes colorectal cancer progression in a CMAH-dependent manner. Specifically, we proposed that a human-like sialic acid glycophenotype characterized by Neu5Ac predominance and Neu5Gc deficiency enhances susceptibility to hyperglycemia, coupling systemic metabolic stress to tumor progression.

To test this hypothesis, we combined a spontaneous colorectal cancer model (CPC-Apc) [24] with streptozotocin (STZ)-induced diabetes [25] in wild-type (WT) and Cmah^-/-^ mice [26], allowing us to define the individual and combined contributions of hyperglycemia and sialome composition to tumor progression and immune remodeling.

## Methods

### Ethics statement

All animal experiments were approved by the Institutional Animal Care and Use Committee of the Federal University of Rio de Janeiro (CEUA/UFRJ; protocol number A06/24-21-21) and conducted in accordance with national and international guidelines. Human tissue samples were obtained under approval of the Institutional Ethics Committee (CEP/Hospital Naval Marcílio Dias; protocol number (39063014.8.0000.5256, Brazil), with informed consent from all participants.

#### Human colorectal cancer samples

Formalin-fixed paraffin-embedded (FFPE) tumor specimens were obtained from patients who underwent surgical resection for colorectal cancer. Based on clinical and laboratory records, patients were stratified into non-diabetic and diabetic groups. Diabetes mellitus was defined according to medical diagnosis. All samples were anonymized before processing and handled in accordance with institutional ethical guidelines. Only specimens containing sufficient tumor area and well-preserved tissue architecture were selected for subsequent histopathological evaluation. Furthermore, samples were matched according to clinical report, tumor stage, age, and anatomical location whenever possible.

**Table 1.** Pairing of human samples according to the adopted criteria.

| Non-diabetic |  |  |  |  | Diabetic |  |  |  |  | Medical report |
| --- | --- | --- | --- | --- | --- | --- | --- | --- | --- | --- |
| ID | Gender | Age | TNM | location | ID | Gender | Age | TNM | location |  |
| 0838/13<sup>4T</sup> | Male | 69 | T4 N1b Mx | SC | 8548/14<sup>7T</sup> | Male | 65 | T3 N1 | DC | moderately differentiated |
| 10659/14<sup>5T</sup> | Male | 49 | T3 N2 | DC | 1586/15<sup>4T</sup> | Female | 67 | T3 N2 | AC/TC | ulcerative and infiltrative well-differentiated |
| 14583/12<sup>7T</sup> | Female | 76 | T3 N0 Mx | SC | 3257/10<sup>3T</sup> | Male | 72 | T3 N0 | AC | moderately differentiated |

All samples were classified as Adenocarcinoma and collected from ascending colon (AC), transverse colon (TC), descending colon (DC) or sigmoid colon (SC).

### Murine model of spontaneous colorectal cancer

*CPC-Apc* WT mice were generated as previously described [24]. To recapitulate the human-specific absence of Neu5Gc, *CPC-Apc* progenitor mice (*Apc*^F/F^ and CDX2P NLS Cre) were crossed with *Cmah*^-/-^ mice, originally described by Hedlund and collaborators [26]. Homozygous deletion of *Cmah* was confirmed by genotyping as described [23], after which *Cmah*^-/-^ progenitors were intercrossed to generate CPC-*Apc Cmah*^-/-^ mice. This genetic background spontaneously develops colon polyps and lack endogenous Neu5Gc synthesis mimicking the human sialic acid repertoire [23]. All animals were maintained with diet free of animal-derived products (irradiated Nuvilab CR-1 chow, cat. no. 276432) to prevent dietary Neu5Gc incorporation.

Animals presenting congenital malformations, severe clinical signs associated with disease progression (including rectal prolapse or anal bleeding), or premature death were excluded according to predefined humane endpoints established by the institutional animal ethics committee.

### Streptozotocin-induced hyperglycemia

Eight-week-old male CPC-*Apc* WT and CPC-*Apc Cmah^-/-^*mice were subjected to 2 hours fasting period prior to baseline measurements of body weight and blood glucose levels. Hyperglycemia was induced using a low-dose (50 mg/kg body weight) streptozotocin (STZ) protocol adapted from Furman [25]. Hyperglycemia was defined as non-fasting blood glucose levels ≥200 mg/dL on two consecutive measurements. Control animals received equivalent volumes of vehicle alone.

All injections were performed at 4:00 p.m. to minimize circadian-related variability in glucose metabolism [27]. Following STZ administration, body weight and blood glucose levels were monitored weekly for 14 weeks. At 5.5 months of age, mice were euthanized by CO₂ inhalation, and tissues were collected for downstream analyses.

### Colonoscopy images acquisition

Longitudinal colonoscopy images were acquired from 8, 14, and 20 weeks-old CPC-*Apc Cmah*^-/-^ mice (corresponding to 1, 6, and 12 weeks after STZ or vehicle treatment) after a previous sedation by isoflurane inhalation, as described by Oliveira and colleagues [28]. In addition, 100 uL [0,02mg/ml] of atropine was injected intraperitoneally to reduce peristalsis.

### Isolation of tumor-infiltrating leukocytes

Tumor-infiltrating leukocytes (TILs) were isolated using an enzymatic digestion and density-gradient protocol optimized for solid tumors. Tumor tissues were excised and mechanically fragmented using a scalpel in 2 mL of serum-free high-glucose Dulbecco’s Modified Eagle Medium (DMEM-HG) containing 0.2% collagenase IV (Sigma-Aldrich, C5138) and 10 µL of DNase I (4mg/mL; RNase-free; New England BioLabs, M0303S). Samples were incubated at 37 °C for 30 min, followed by gentle mechanical dissociation using a glass Pasteur pipette and a second incubation for an additional 30 min at 37 °C.

Enzymatic digestion was stopped by addition of EDTA (0.5 M; 1:100 dilution), and cell suspensions were filtered through a nylon mesh into 15 mL tubes and centrifuged at 400g for 5 min. Pellets were resuspended in 100 µL of FACS buffer (PBS supplemented with 2% fetal bovine serum), transferred to a “u” bottom plate and centrifuged for subsequent flow cytometry staining and analysis.

### Flow cytometry analysis

Fresh cells were pre-incubated with anti-CD16/32 (Fc block) and then proceeded to the proposed staining (antibodies list available in supplementary material) for 20 min at 4 °C using a 1:200 dilution, except for PE-conjugated antibodies, which were used at a 1:400 dilution. Following staining, cells were centrifuged at 400g for 5 min, washed with FACS buffer and acquired on a BD LSRFortessa™ X-20 flow cytometer. Data analysis was performed using FlowJo^®^ software.

### Histopathological analysis

Formalin-fixed, paraffin-embedded sections (5 μm) from murine colons and human colorectal tumors were stained with hematoxylin and eosin (H&E) and digitized using whole-slide scanners. Histopathological evaluation was performed by a pathologist blinded to the experimental groups. Dysplastic lesions were classified as low-or high-grade according to established morphological criteria. The stromal compartment was quantified by manually delineating the mesenchymal area on digital whole-slide images using ImageJ (NIH) and expressing it as a percentage of the total tumor area.

### Statistical analysis

Statistical analyses were performed using GraphPad Prism 10. Normality was assessed using the Shapiro–Wilk test prior to parametric analyses. Comparisons between two groups were performed using Student’s *t*-test or Mann–Whitney test, as appropriate. Multiple group comparisons were analyzed by one-way ANOVA followed by Tukey post hoc test or two-way ANOVA, where applicable. Biological replicates (individual animals) were used for all analysis. Adjustment for multiple comparisons was applied where appropriate.

## Results

### Reproduction of the reduced spontaneous tumorigenesis phenotype in CPC*-Apc Cmah^-/-^* mice

As a prerequisite for investigating the impact of hyperglycemia, we first reproduced the CPC-*Apc Cmah^-/-^* model previously established by our group, which combines colon-specific *Apc* inactivation with constitutive *Cmah* deficiency to recapitulate the human-specific absence of endogenous Neu5Gc (Fig. 1A). Consistent with our previous report [23], macroscopic analysis revealed marked differences in tumor burden between genotypes. CPC-*Apc* WT mice developed numerous large colonic polyps, whereas CPC-*Apc Cmah*^-/-^ animals exhibited only small, sporadic lesions consistent with a milder disease phenotype (Fig. 1B). Quantitative analyses confirmed significant reductions in both polyp number (Fig. 1C) and tumor burden index, calculated as the cumulative diameter of all polyps per mouse (Fig. 1D). These findings validate the previously reported phenotype in our experimental cohort and establish a robust baseline for assessing the effects of STZ-induced hyperglycemia.

**Fig. 1.**
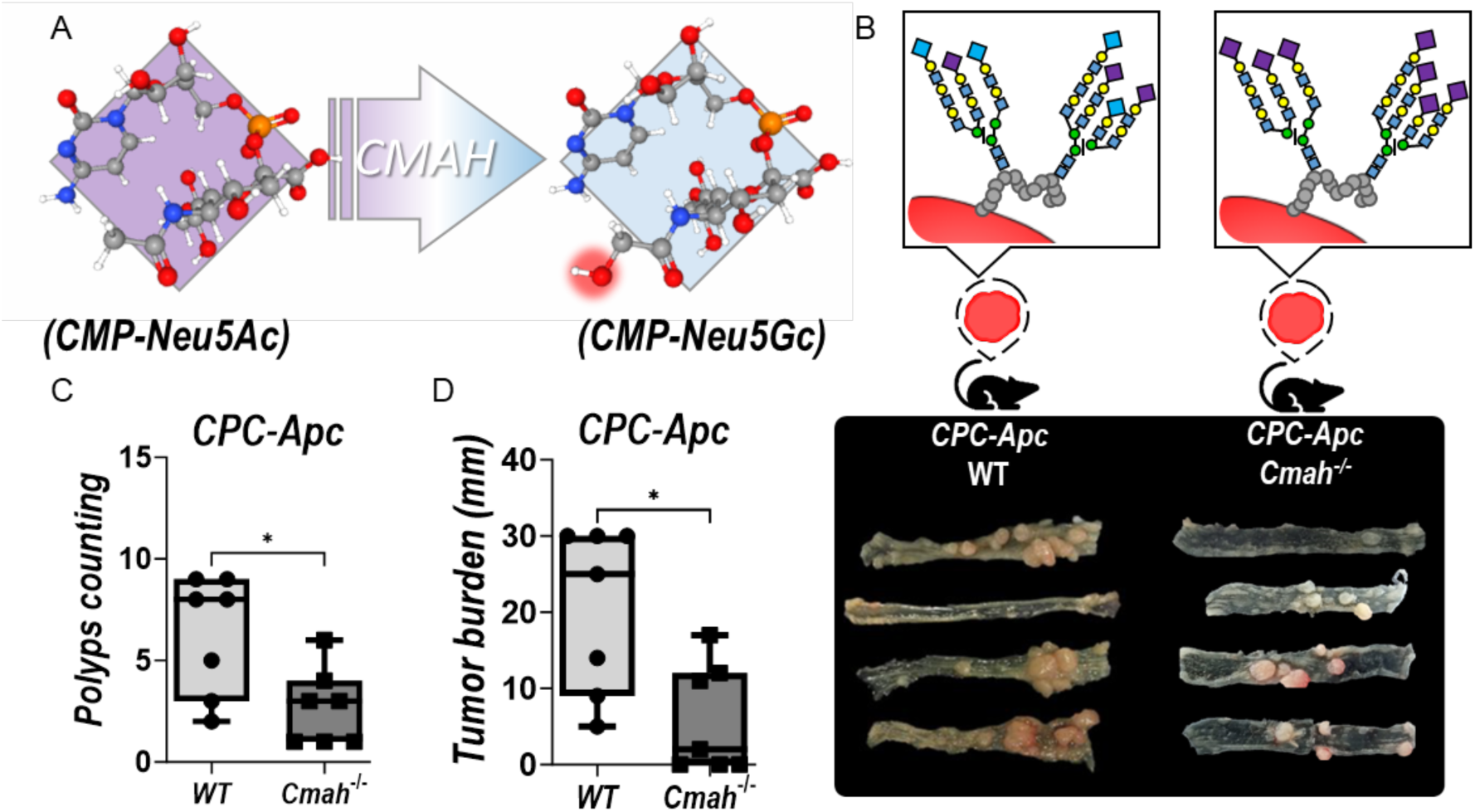
Cmah deficiency attenuates spontaneous colorectal tumorigenesis. (A) Schematic representation of CMAH enzymatic activity converting CMP-Neu5Ac (left, PubChem CID: 448209) to CMP-Neu5Gc (right; PubChem CID: 126897); (B) Upper panels: Representative schematic models of cell surface sialylated glycans in CPC-Apc WT and *Cmah^-/-^* mice. Loss of *Cmah* abolishes Neu5Gc biosynthesis, leading to exclusive incorporation of Neu5Ac into cell surface glycoconjugates. Lower panels: Representative images of colons from 5.5-month-old CPC-*Apc* WT and CPC-*Apc Cmah^-/-^* mice showing spontaneous colorectal tumor development. (C) Quantification of the number of polyps per animal; (D) Tumor burden index calculated as the sum of polyp diameters per mouse. Data are presented as mean ± SEM. Statistical significance was determined using an unpaired two-tailed Student’s *t*-test; *p* < 0.05. n_WT_ = 7; n_Cmah-/-_ = 7.

### CMAH deficiency reshapes the tumor-infiltrating leukocyte landscape

Having confirmed the reduced spontaneous tumor burden in CPC-*Apc Cmah^-/-^* mice, we next asked whether this phenotype was associated with differences in antitumor immunity, since the composition of the tumor immune infiltrate is a major determinant of colorectal cancer progression and clinical outcome [29]. To address this question, we analyzed tumor-infiltrating leukocyte (TIL) populations in polyps from CPC-*Apc* WT and CPC-*Apc Cmah^-/-^* mice by flow cytometry as described in Supplementary figure 1. Although CPC-*Apc Cmah^-/-^*mice exhibited a notable increase in CD45^+^ cells (Fig. 2A), we observed distinct distributions of specific leukocyte populations. First, we found a significant reduction in Natural Killer (NK) cells (NK1.1⁺ CD3⁻), and no differences in NKT cells (NK1.1⁺CD3⁺) (Fig. 2B–C). Moreover, CPC-*Apc Cmah^-/-^* mice displayed a marked increase in intratumoral B cells (CD19⁺CD3⁻), accompanied by reduced infiltration of T cells (CD19⁻CD3⁺), and no significant differences in CD4^+^ or CD8^+^ T-cell subsets (Fig. 2D–G). In addition, the overall frequency of tumor-associated macrophages (TAMs; CD11b⁺F4/80⁺) was significantly lower in CPC-*Apc Cmah*^-/-^ mice compared with WT controls. However, both genotypes displayed a similar distribution of M1-like (CD86⁺CD206⁻), mixed (CD86⁺CD206⁺), and M2-like (CD86⁻CD206⁺) macrophage subsets (Fig. 2H–K).

**Fig. 2.**
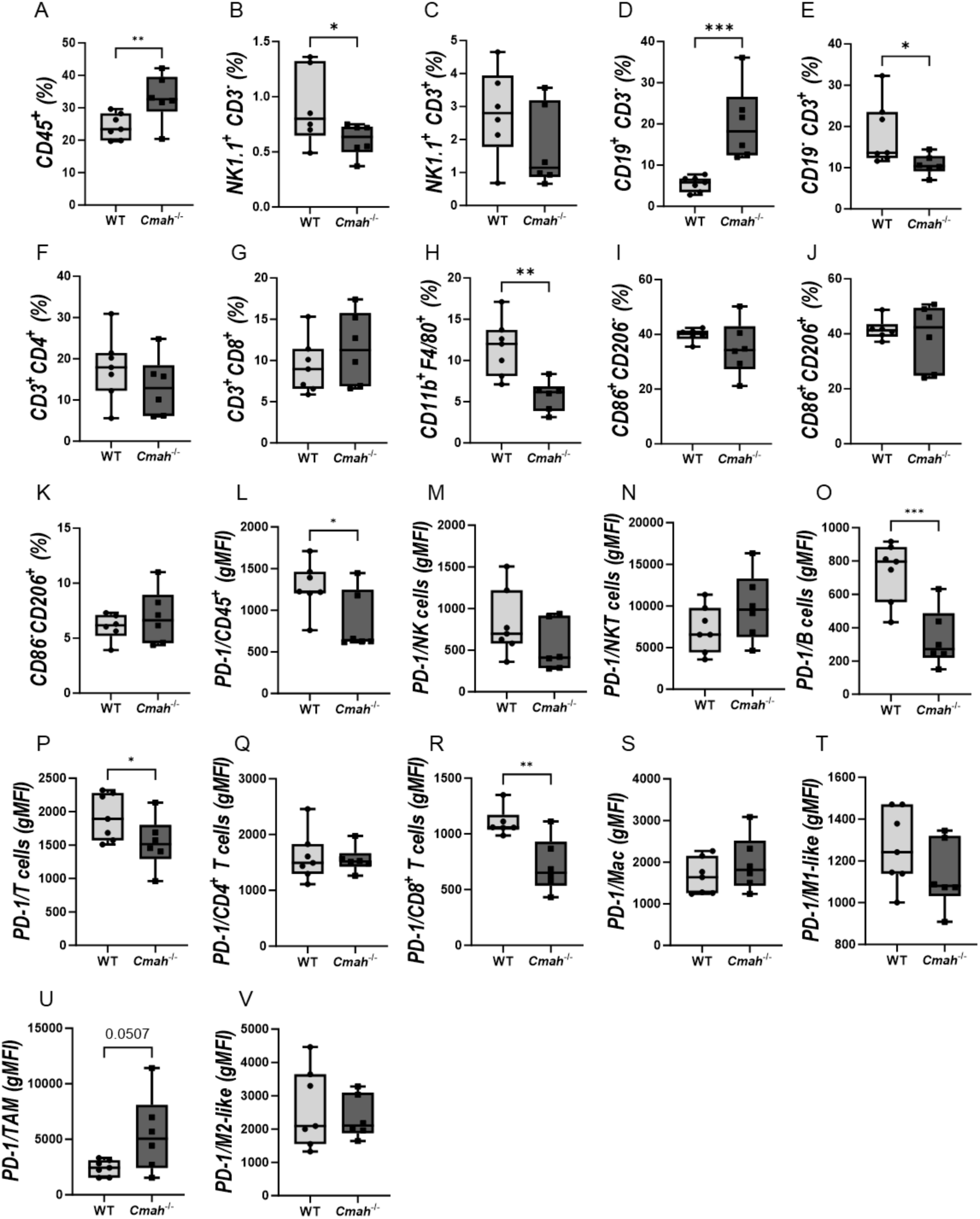
CMAH deficiency reshapes the tumor-infiltrating leukocyte landscape. (A) Total TILs frequency; (B-G) Frequencies of lymphoid-derived immune cell populations. (H) Total tumor-associated macrophages (TAMs; CD11b⁺F4/80⁺). (I-K) Distribution of TAM subsets: M1-like (CD86⁺CD206⁻), mixed (CD86⁺CD206⁺), and M2-like (CD86⁻CD206⁺). (L-V) Comparison of Geometric MFI of PD-1 in all populations analyzed. Data are presented as mean ± SEM. Statistical significance was determined using an unpaired two-tailed Student’s t-test or Mann– Whitney test, as appropriate. p < 0.05; n_WT_ = 7 and n_Cmah–/–_ = 6.

To investigate the exhaustion status of TILs, we analyzed the surface expression of programmed death protein 1 (PD-1), an inhibitory receptor associated with tumor immune evasion [30]. We initially observed a significant reduction in PD-1 expression among total CD45^+^ cells, suggesting a reduced PD-1 expression in CPC-*Apc Cmah^-/^*^-^ mice (Fig. 2L). When individual populations were analyzed, we identified a marked reduction in PD-1 expression in B cells and cytotoxic CD8^+^ T cells from CPC-*Apc Cmah^-/^*^-^ mice compared with CPC-*Apc* WT animals (Fig. 2M–R). Conversely, no significant changes in PD-1 expression were detected among TAM populations (Fig. 2S–V).

Together, these findings demonstrate that CMAH deficiency leads to substantial reshape of the immune composition of the tumor microenvironment (TME), enrichment of intratumoral PD-1^low^ B cells, and accumulation of reduced PD-1 expressing cytotoxic CD8^+^ T cells. These results suggest that human-specific sialic acid patterns influence immune cell composition and activation status within the tumor microenvironment.

### STZ-induced hyperglycemia is consistently established in both CPC-Apc genotypes

To establish a hyperglycemic background, CPC-*Apc* WT and CPC-*Apc Cmah*^-/-^ mice were treated with low-dose streptozotocin (STZ) (Fig. 3A). STZ induced body weight loss and sustained hyperglycemia in both genotypes (Fig. 3B–E). Although CPC-*Apc Cmah*^-/-^ mice displayed slightly lower basal glycemia than WT controls, both genotypes reached comparable hyperglycemic levels following STZ treatment (Fig. 3F), validating the model for subsequent analyses of diabetes-associated colorectal cancer progression.

**Fig. 3.**
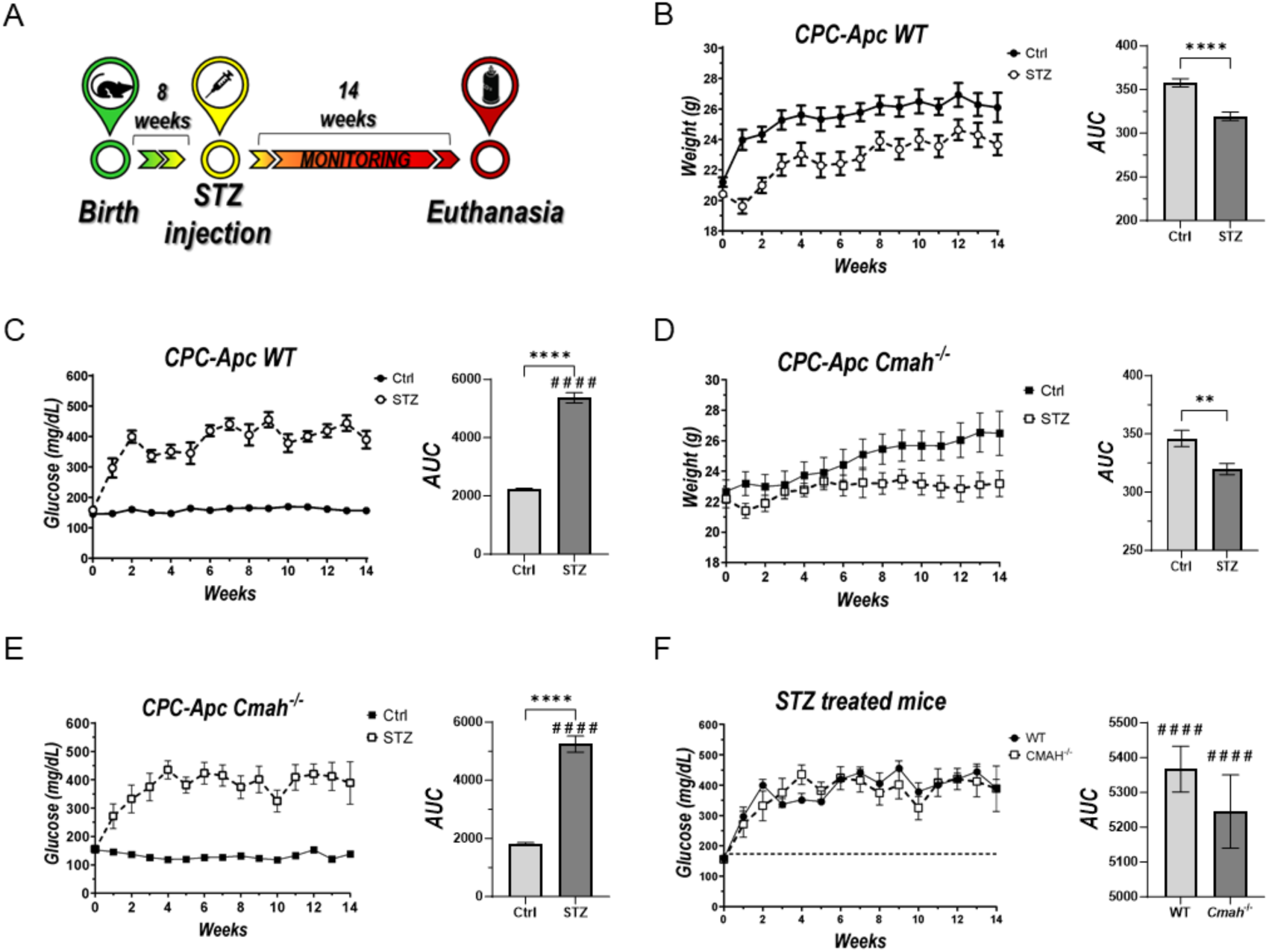
STZ-induced hyperglycemia is consistently established in CPC-*Apc* WT and *Cmah^-**/-**^* mice. (A) Experimental timeline of the low-dose streptozotocin (STZ)-induced hyperglycemia protocol. (B and C) Body weight and blood glucose monitoring in CPC-*Apc* WT mice throughout the experimental period. (D and E) Body weight and blood glucose monitoring in CPC-*Apc Cmah*^-/-^ mice throughout the experimental period. (F) Comparison of hyperglycemic status induced by STZ treatment between genotypes. Data are presented as mean ± SEM. A one-sample Student’s t-test was performed to compare blood glucose levels in STZ-treated mice against a hypothetical threshold value of 200 mg/dL (**####** p < 0.0001). Comparisons of area under the curve (AUC) values were performed using an unpaired two-tailed Student’s t-test (** p < 0.01; **** p < 0.0001). nn_WTctrl_= 7, n_WTstz_= 7, n_CMAHctrl_^-/-^= 7 and n_CMAHstz_^-/-^= 7.

### Hyperglycemia exacerbates CRC progression in CPC-Apc Cmah^-/-^ mice

Having established comparable hyperglycemia in both genotypes, we next investigated its impact on colorectal tumor progression. Macroscopic analysis revealed that STZ treatment did not significantly alter either the number or morphology of colonic polyps in WT animals (Fig. 4A). In striking contrast, CPC-*Apc Cmah^-/-^*mice exhibited a pronounced worsening of disease severity under hyperglycemia.

**Fig. 4.**
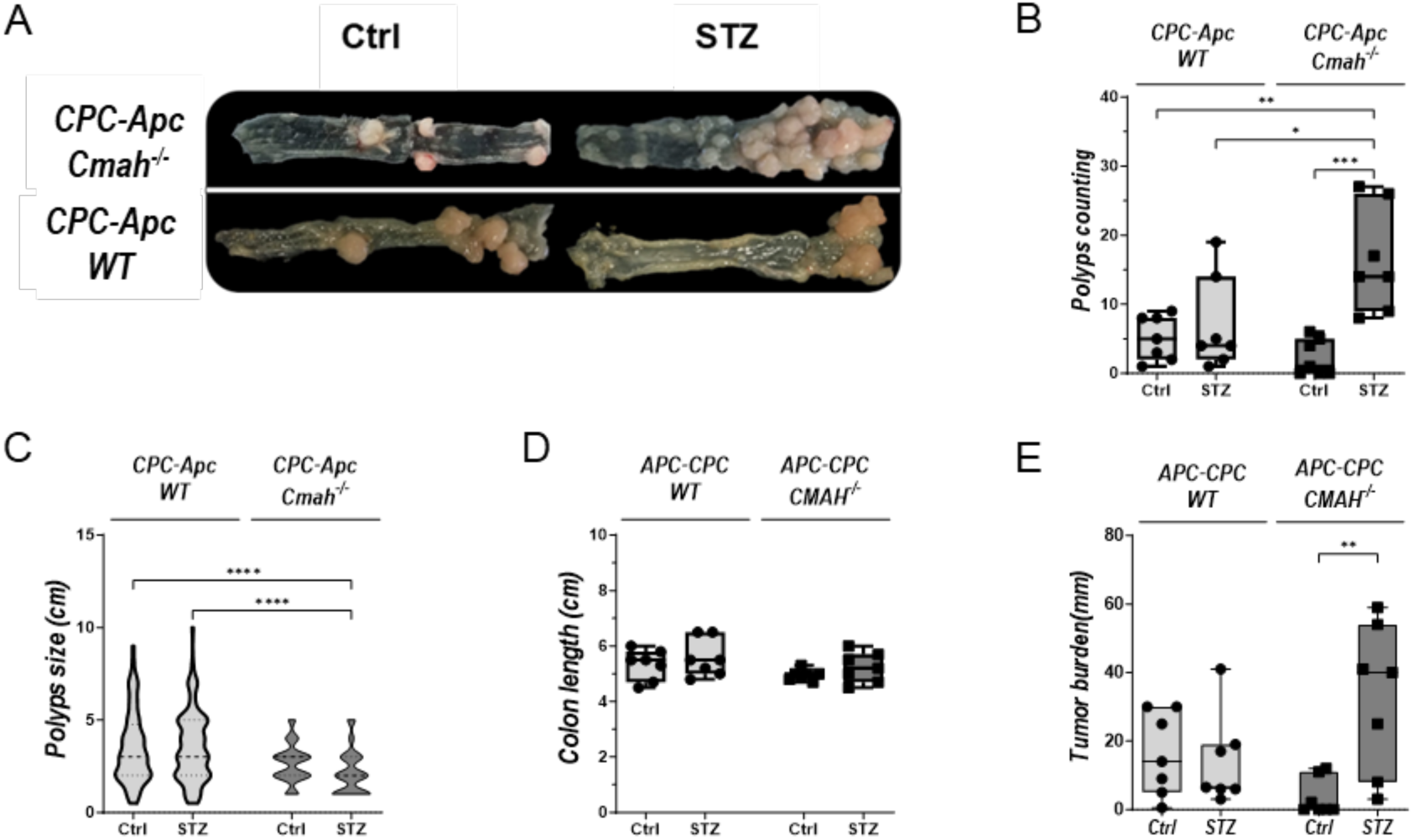
Hyperglycemia selectively exacerbates colorectal cancer progression in CPC-*Apc Cmah*^-/-^ mice. (A) Representative images of colons from CPC-*Apc* WT and CPC-*Apc Cmah*^−/−^ mice following vehicle (Ctrl) or STZ treatment, showing spontaneous colorectal tumor formation. (B) Number of polyps per animal. (C) Individual polyps size distribution. (D) Colon length measurements. (E) Tumor burden index calculated as the sum of polyps diameters per mouse. Data are presented as mean ± SEM. Statistical analyses were performed using ordinary two-way ANOVA. Significance levels are indicated as follows: *p* < 0.0332 (*)*, p < 0.0021 (\*\**)*, p < 0.0002 (\*\*\**), and *p* < 0.0001 (****). n_WTctrl_= 7, n_WTstz_= 7, n_CMAHctrl_^-/-^= 7 e n_CMAHstz_^-/-^= 7.

Hyperglycemic CPC-*Apc Cmah*^-/-^ mice developed substantially more colonic lesions than their euglycemic counterparts, as evidenced by gross inspection of the colon (Fig. 4A). Quantitative analysis confirmed that, while polyp numbers remained unchanged in CPC-*Apc* WT mice following STZ treatment, hyperglycemia induced an approximately fourfold increase in polyp number in hyperglycemic CPC-*Apc Cmah*^-/-^ mice compared with CPC-*Apc Cmah*^-/-^ controls (Fig. 4B). Moreover, hyperglycemic CPC-*Apc Cmah*^-/-^ mice displayed significantly higher polyp counts than both WT groups, including hyperglycemic CPC-*Apc* WT animals, highlighting a genotype-related susceptibility to the tumor-promoting effects of hyperglycemia.

Analysis of individual polyp size revealed no significant difference between STZ-treated and control CPC-*Apc Cmah*^-/-^ mice. However, polyps arising in CPC-*Apc Cmah*^-/-^ mice were significantly smaller than those observed in CPC-*Apc* WT animals, irrespective of glycemic status (Fig. 4C). Colon length did not differ among experimental groups (Fig. 4D), indicating that the observed differences were not attributable to gross anatomical alterations. Importantly, despite similar polyp sizes, the cumulative tumor burden, calculated as the sum of all polyp diameters per mouse, was markedly increased in STZ-treated CPC-*Apc Cmah*^-/-^ mice relative to all WT groups (Fig. 4E).

Together, these results demonstrate that CPC-*Apc Cmah^-/-^* mice showed more susceptibility to the tumor-promoting effects of hyperglycemia. This phenotype mirrors the poorer clinical outcomes observed in diabetic CRC patients and supports the concept that absence of Neu5Gc worsens tumor progression under glucose-induced metabolic stress.

### Hyperglycemia accelerates the progression of colorectal lesions in Cmah^-/-^ mice

Given the increased tumor burden observed in hyperglycemic CPC-*Apc Cmah^-/-^* mice, we next asked whether hyperglycemia also accelerated the temporal development of colorectal lesions. Longitudinal colonoscopy was performed in CPC-*Apc Cmah^-/^*^-^ mice at 8, 14, and 20 weeks of age (corresponding to 1, 6, and 12 weeks after STZ or vehicle treatment), enabling *in vivo* monitoring of polyp growth over time.

At week 8, no visible polyps were detected in either group. However, shortly after STZ administration, small mucosal protrusions suggestive of early lesions were observed only in hyperglycemic mice (Fig. 5, left panel). By week 14, hyperglycemic CPC-*Apc Cmah*^-/-^ mice displayed well-developed colonic polyps, whereas euglycemic animals exhibited only minor mucosal alterations (Fig. 5, middle panel). At the final time point (week 20), hyperglycemic mice showed extensive polyp growth and near-complete luminal occlusion, contrasting with the milder phenotype observed in euglycemic controls (Fig. 5, right panel).

**Fig. 5.**
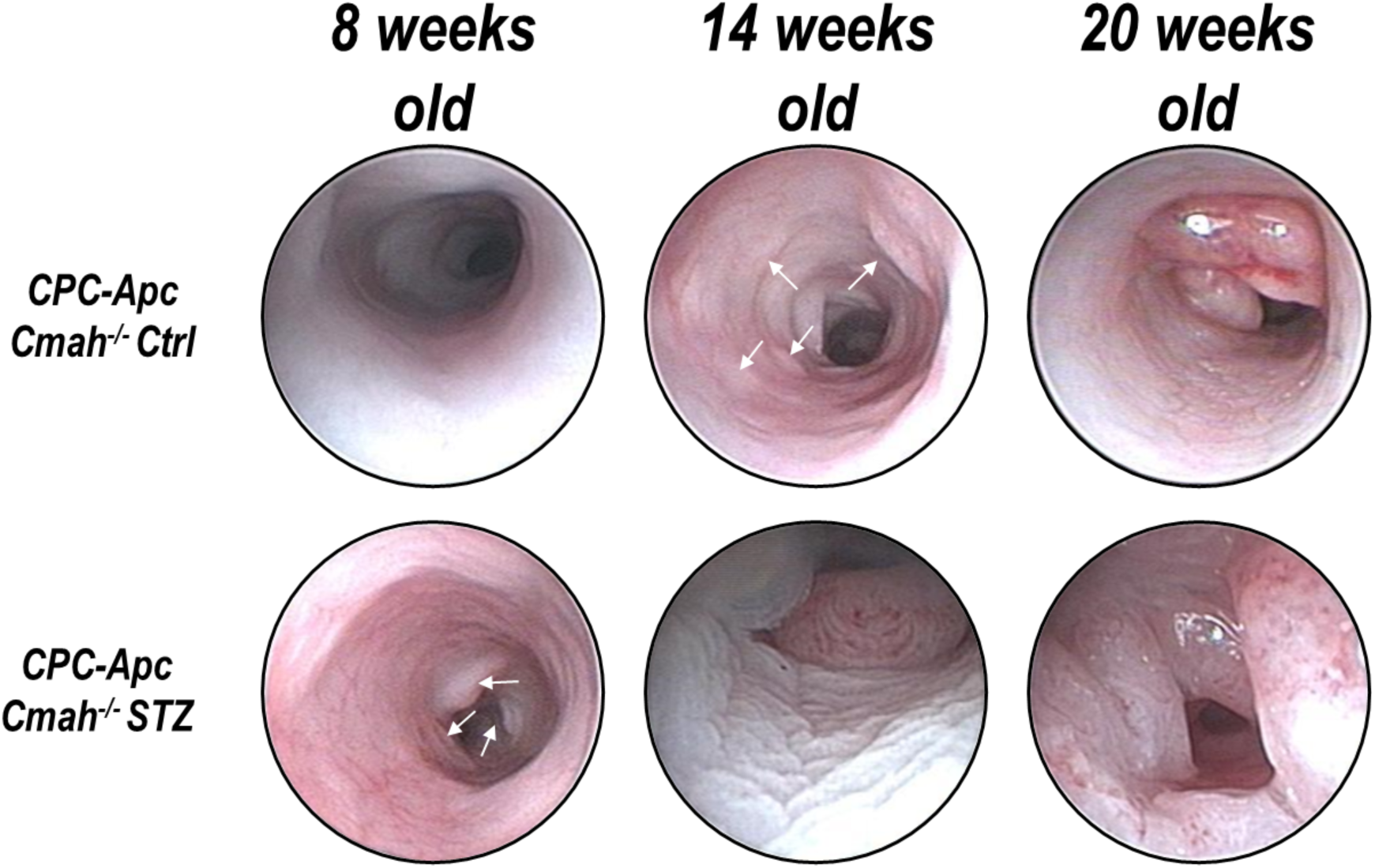
Representative colonoscopic images of colorectal lesion development following STZ treatment in CPC-*Apc Cmah*^-/-^ mice. Representative colonoscopic images acquired at 8, 14, and 20 weeks of age (corresponding to 1, 6, and 12 weeks after treatment) following vehicle or STZ treatment, illustrating the temporal progression of colonic alterations. White arrows indicate suspected early lesions detected during longitudinal monitoring.

Together, these findings demonstrate that STZ-induced hyperglycemia accelerates the onset and progression of colorectal lesions in CPC-*Apc Cmah*^-/-^ mice, reinforcing the genotype-related susceptibility to hyperglycemia-induced worsening observed in tumor burden analysis.

### CPC-Apc Cmah^-/-^ mice recapitulate the histopathological features of human colorectal cancer

To determine whether CMAH deficiency influences tumor malignancy, we analyzed dysplastic lesions and tissue architecture in murine and human colorectal cancer samples.

Under normoglycemic conditions, tumors from CPC-*Apc* WT displayed marked architectural disruption, including hyperchromasia, crypt collapse, and loss of epithelial organization. In contrast, polyps from euglycemic CPC-*Apc Cmah*^-/-^ retained partially preserved tissue architecture and features consistent with low-grade dysplasia (Fig. 6A). Following STZ-induced hyperglycemia, however, CPC-*Apc Cmah*^-/-^ mice developed tumors with a substantially more aggressive histological profile resembling that observed in WT animals.

**Fig. 6.**
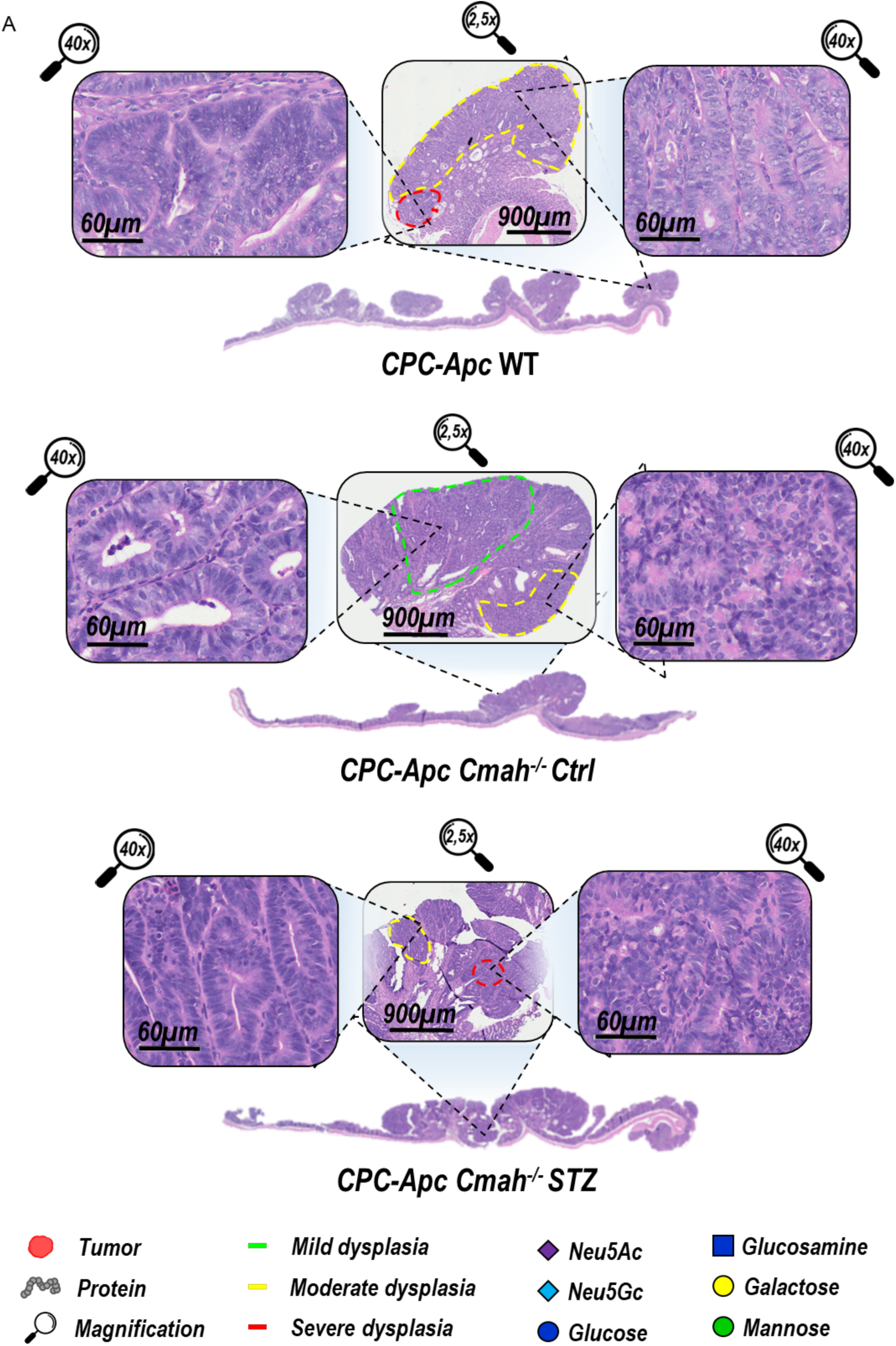
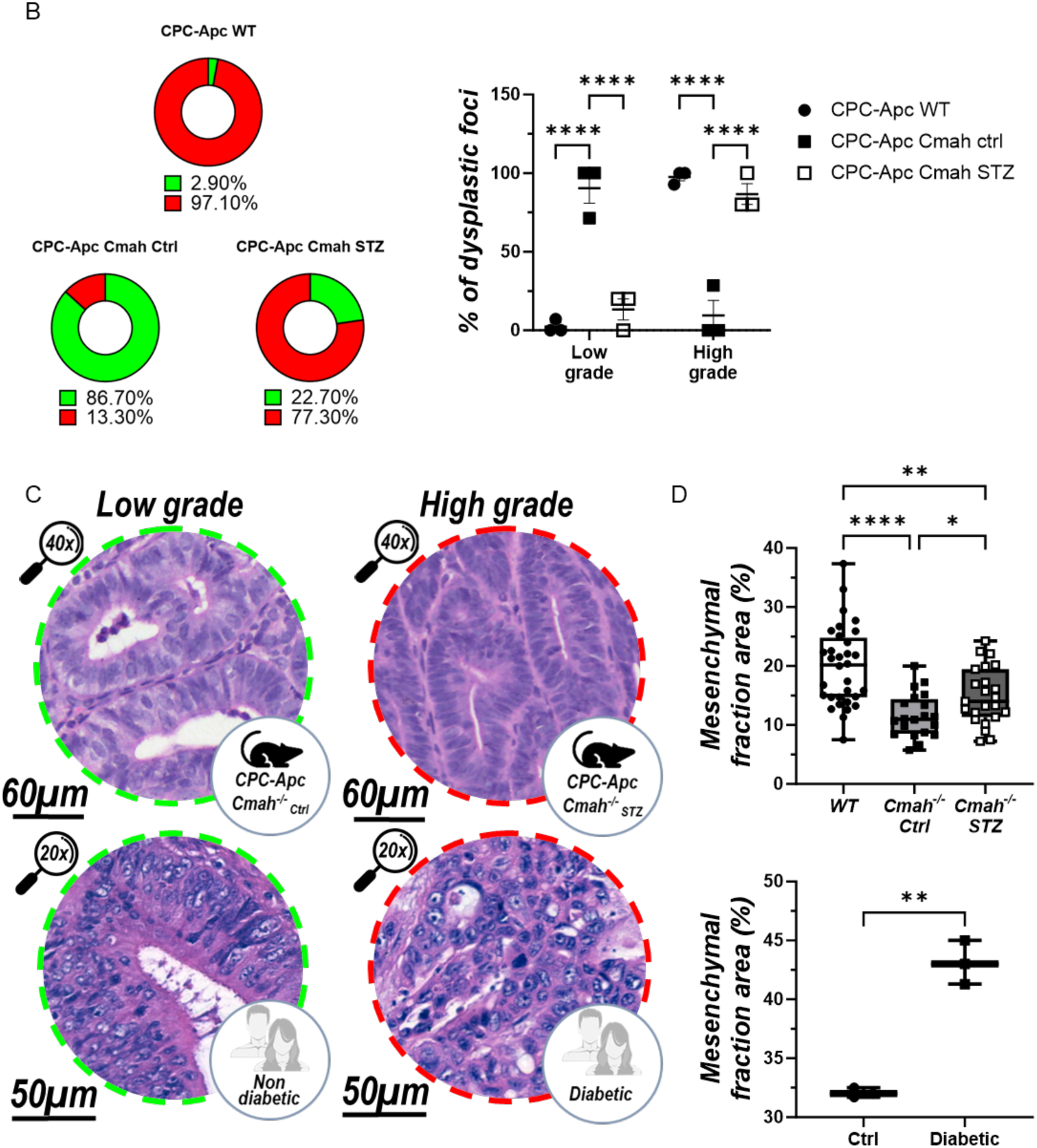
Histopathological alterations associated with Cmah deficiency and hyperglycemia in murine and human colorectal cancer. (A) Representative H&E-stained colorectal tumors sections from CPC-*Apc* WT, CPC-*Apc Cmah*^-/-^ and CPC-*Apc Cmah*^-/-^ STZ treated mice. Insets show representative low- and high-magnification images highlighting tumor architecture and dysplastic regions. Colored outlines indicate histopathological regions classified according to dysplasia grade. (B) Quantitative distribution of dysplastic lesions classified as low-grade (green) or high-grade (red) among experimental groups. (C) Representative histological features of colorectal tumors from the murine model (upper panels) and human colorectal cancer samples (lower panels), comparing control/non-diabetic and hyperglycemic/diabetic conditions. (D) Quantification of mesenchymal area relative to total tumor area in murine (upper panel) and human (lower panel) colorectal tumor samples. Data are presented as mean ± SEM. Statistical analyses for murine samples were performed using Brown–Forsythe and Welch ANOVA tests \**p* < 0.05, \*\**p* < 0.01, \*\*\*\**p < 0.0001. Human sample comparisons were performed using an unpaired Student’s t-test with Welch’s correction (p < 0.01).* n_WTctrl_= 3, n_CMAHctrl_^-/-^= 3, n_CMAHstz_^-/-^= 3, n_non-diabetic_ = 3.

Quantitative grading confirmed reduced disease severity in euglycemic CPC-*Apc Cmah*^-/-^ mice, whereas hyperglycemic CPC-*Apc Cmah*^-/-^ animals showed a marked shift toward high-grade dysplasia (Fig. 6B). Importantly, tumors from CPC-*Apc Cmah*^-/-^ mice closely resembled colorectal lesions from non-diabetic and diabetic patients under normoglycemic and hyperglycemic conditions, respectively (Fig. 6C; supplementary Fig. 2).

Consistently, tumors from euglycemic CPC-*Apc Cmah*^-/-^ exhibited reduced mesenchymal content compared with WT tumors, whereas hyperglycemic CPC-*Apc Cmah*^-/-^ mice developed expanded stromal compartments similar to those observed in tumors from diabetic CRC patients (Fig. 6D).

Together, these findings demonstrate that CPC-*Apc Cmah*^-/-^ mice recapitulate key histopathological features of human colorectal cancer under both normoglycemic and hyperglycemic conditions, reinforcing the translational relevance of this model for studying diabetes-associated colorectal carcinogenesis.

## Discussion

Approximately three million years ago, the evolutionary inactivation of the *CMAH* gene eliminated endogenous Neu5Gc synthesis in the human lineage, establishing a sialic acid repertoire dominated by Neu5Ac [18]. This singular event has been linked to both enhanced immune reactivity [19, 20] and an increased predisposition to metabolic disturbances [21]. More recently, perturbations in the Neu5Ac/Neu5Gc balance, whether through transcriptional reactivation of the *CMAH* pseudogene [22] or dietary incorporation of Neu5Gc [23], have been associated with heightened aggressiveness of gastrointestinal tumors. We reasoned that the distinct sialome imposed by CMAH loss may reshape the tumor microenvironment in ways that alter its susceptibility to exogenous metabolic insults. Building upon this framework, we hypothesized that the human-like sialome regulates the colorectal tumor microenvironment, rendering it selectively vulnerable to the pro-tumor effects of chronic hyperglycemia.

Our findings directly support this hypothesis. Under euglycemic conditions, CPC-*Apc Cmah^-/-^* mice exhibited reduced spontaneous colorectal tumorigenesis compared with WT counterparts, since that endogenous Neu5Gc may facilitate proliferative signaling or tumor–stroma communication [22, 23]. This protective phenotype was accompanied by substantial remodeling of the tumor immune microenvironment, including increased recruitment of tumor-infiltrating leukocytes and enrichment of B lymphocytes. Although antitumor immunity in CRC has been predominantly studied in the context of T-cell responses, emerging evidence suggests that tumor-infiltrating B cells can exert important immunoregulatory functions within the tumor microenvironment [31–33]. Interestingly, reduced PD-1 expression was observed predominantly in B cells and cytotoxic CD8⁺ T cells, is consistent with a bias against exhaustion within these populations.

Since Diabetes-derived hyperglycemia is considered a critical “second hit” in tumor progression [5, 11, 34–37], we next examined its impact in this model. Although STZ-induced hyperglycemia elevated blood glucose to comparable levels in both CPC-*Apc* WT and *Cmah*^-/-^ mice, hyperglycemia exacerbated tumor progression exclusively in the *Cmah*^-/-^ background. This genotype-specific susceptibility was evident in the increased polyp number and tumor burden, the accelerated temporal onset of lesions visualized by longitudinal colonoscopy, and the histopathological shift toward high-grade dysplasia with expanded mesenchymal components [38]. The resemblance between tumors from hyperglycemic CPC-*Apc Cmah*^-/-^ mice and those from diabetic CRC patients underscores the translational relevance of this model, positioning it as a valuable model to study diabetes-associated colorectal carcinogenesis.

While our study establishes a robust phenotypic link between CMAH deficiency, hyperglycemia, and accelerated tumor progression, the underlying molecular mechanisms remain to be elucidated. One possible mechanism involves hypersialylation of surface glycoconjugates [9, 11, 15]. Such remodeling of the tumor glycocalyx could alter glycan-mediated immune recognition and favor engagement of inhibitory Siglec receptors, particularly Siglec-9 and Siglec-10, which have been implicated in colorectal cancer-associated immune suppression and tumor immune escape [39–41]. In parallel, hyperglycemia may synergize with oncogenic pathways such as Wnt, MAPK, and PI3K through enhanced O-GlcNAcylation [9, 11, 12], thereby counteracting the baseline tumor-suppressive phenotype observed in CPC-*Apc Cmah^-/^*^-^ mice. Future studies integrating glycoproteomics, metabolic tracing, and functional Siglec blockade can identify the potencial molecular interplay between altered sialylation, metabolic rewiring, and immune regulation in this context.

An important aspect of our experimental design is that the STZ model primarily recapitulates hyperglycemia associated with relative insulin deficiency, rather than the hyperinsulinemic state characteristic of type 2 diabetes [42, 43]. This distinction partially isolates the glycemic component from other metabolic alterations linked to diabetes-associated CRC. Importantly, STZ treatment alone did not exacerbate tumor progression in WT mice, supporting the interpretation that the observed phenotype results from the interplay between hyperglycemia and the Neu5Ac-dominant sialome rather than nonspecific STZ toxicity. Although multiple mechanisms contribute to diabetes-associated CRC progression, including chronic inflammation and hyperinsulinemia [44], our findings indicate that hyperglycemia alone is sufficient to potentiate tumor progression in a CMAH-dependent context. Therefore, the protumoral effects observed in our model may underestimate the full oncogenic burden experienced by diabetic patients, further reinforcing the biological relevance of the phenotype identified in this study.

Some limitations of the present study should be acknowledged. Although the model captures the absence of endogenous Neu5Gc characteristic of humans, it does not fully reproduce the complexity of the human glycoimmune system. In addition, direct glycomic profiling, microbiome characterization, and functional validation of immune subsets were beyond the scope of the current work and should be addressed in future studies.

In summary, our findings demonstrate that the human-specific sialome established by the evolutionary loss of CMAH is not merely a passive evolutionary trait but a determinant of colorectal cancer susceptibility under hyperglycemia-induced metabolic stress. By reproducing this human-specific glycosylation phenotype, the CPC-*Apc Cmah^-/^*^-^ model provides a unique translational platform to investigate the mechanisms linking diabetes to colorectal carcinogenesis. More broadly, our findings suggest that the evolutionary loss of CMAH, while defining a hallmark of human glycan biology, may have inadvertently increased human susceptibility to hyperglycemia-associated colorectal cancer.

## Acknowledgements

This work was supported by funding from Coordenação de Aperfeiçoamento de Pessoal de Nível Superior, Conselho Nacional de Desenvolvimento Científico e Tecnológico and Fundação de Amparo à Pesquisa do Estado do Rio de Janeiro. We also thank the multi-users platforms from UFRJ: Plataforma de Imunoanálise (PIA), Plataforma de Microscopia Óptica de Luz Gustavo de Oliveira Castro (PLAMOL), Plataforma de Modelos Biológicos Fernando Garcia de Mello (PLAMB) and Unidade Multiusuário de Citometria. Artificial intelligence–assisted technology (ChatGPT, OpenAI, San Francisco, CA, USA) was used solely for language refinement and grammatical editing of the manuscript. All scientific content, interpretation, and final revisions were performed and validated by the authors.

## Credit authorship contribution statement

**Ronan C. M. Santos**: Writing – review & editing, Writing – original draft, Visualization, Validation, Methodology, Investigation, Formal analysis, Data curation, Conceptualization. **Giulia S. Ferreira**: Writing – review & editing, Methodology, Investigation, Formal analysis, Data curation. **Agata C. Fonseca**: Methodology, Validation, Formal analysis. **Cristina Maeda Takiya**: Writing – review & editing, Methodology, Validation, Formal analysis, Data curation. **Ana Luiza Lopes**: Review & editing, Methodology, Validation, Data curation. **Miguel Fontes**: Methodology, Validation, Formal analysis, Data curation. **César S. Bastos Junior**: Resources, Methodology. **Vanessa H. N. da Silva**: Methodology, Formal analysis, Data curation. **João C. Machado**: Resources, Methodology, Formal analysis, Data curation. **Miriam B. F. Werneck**: Writing – review & editing, Supervision, Formal analysis, Data curation. **Wagner B. Dias**: Writing – review & editing, Resources, Funding acquisition. **Frederico Alisson-Silva**: Writing – review & editing, Supervision, Resources, Methodology, Conceptualization. **Adriane Regina Todeschini**: Writing – review & editing, Supervision, Resources, Project administration, Methodology, Funding acquisition, Conceptualization.

## SUPPLEMENTARY MATERIAL

**Supplementary table 1.** Antibodies list for Flow cytometry analysis.

| Marker | Fluorophore | Catalog number |
| --- | --- | --- |
| CD45 | PercP Cy5.5 | eBioscience, 45-0451-82 |
| NK1.1 | BV421 | BioLegend, 108732 |
| CD19 | APC Cy7 | BioLegend, 115530 |
| CD3ε | FITC | BioLegend, 155603 |
| CD4 | BV605 | BD Biosciences, 563151 |
| CD8 | PE | ImmunoTools, 22150084sp |
| F4/80 | BV421 | BioLegend, 123132 |
| CD11b | FITC | BioLegend, 101206 |
| CD86 | BV605 | BioLegend, 105037 |
| CD206 | PE | BioLegend, 141706 |
| CD279 (PD-1) | PE Cy7 | Biolegend, 135215 |
| CD16/CD32<br>(Fc block) | Purified | BD Pharmigen, 553142 |

**Supplementary Fig. S1.**
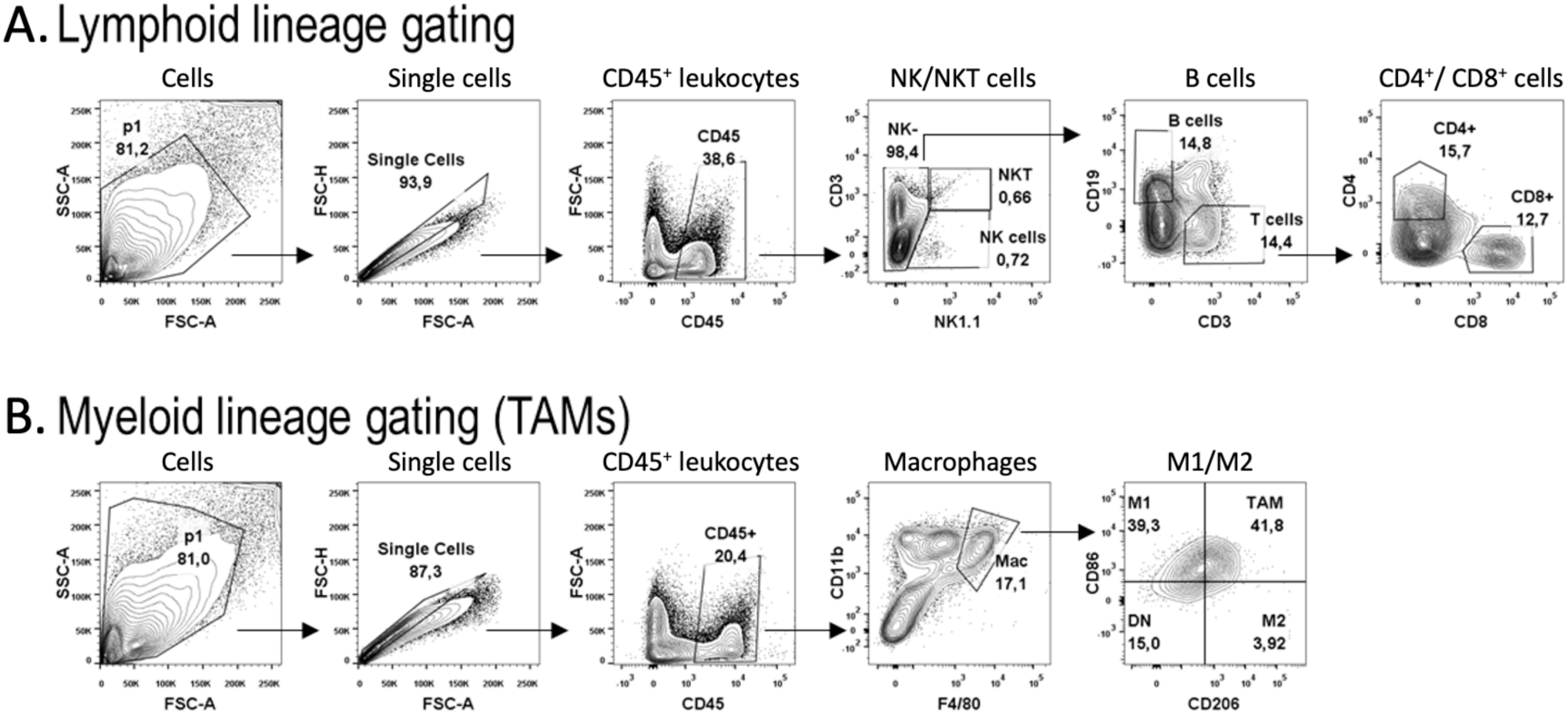
Flow cytometry gating strategy used to identify tumor-infiltrating leukocyte populations. Sequential gating included exclusion of debris, doublets, and dead cells, followed by identification of CD45⁺ leukocytes. Lymphoid populations (A) were further subdivided into NK, NKT, B, CD4⁺ and CD8⁺ T cells, whereas myeloid cells (B) were classified as tumor-associated macrophages (TAMs) and subsequently separated into M1-like, M2-like and mixed populations.

**Supplementary Fig. S2.**
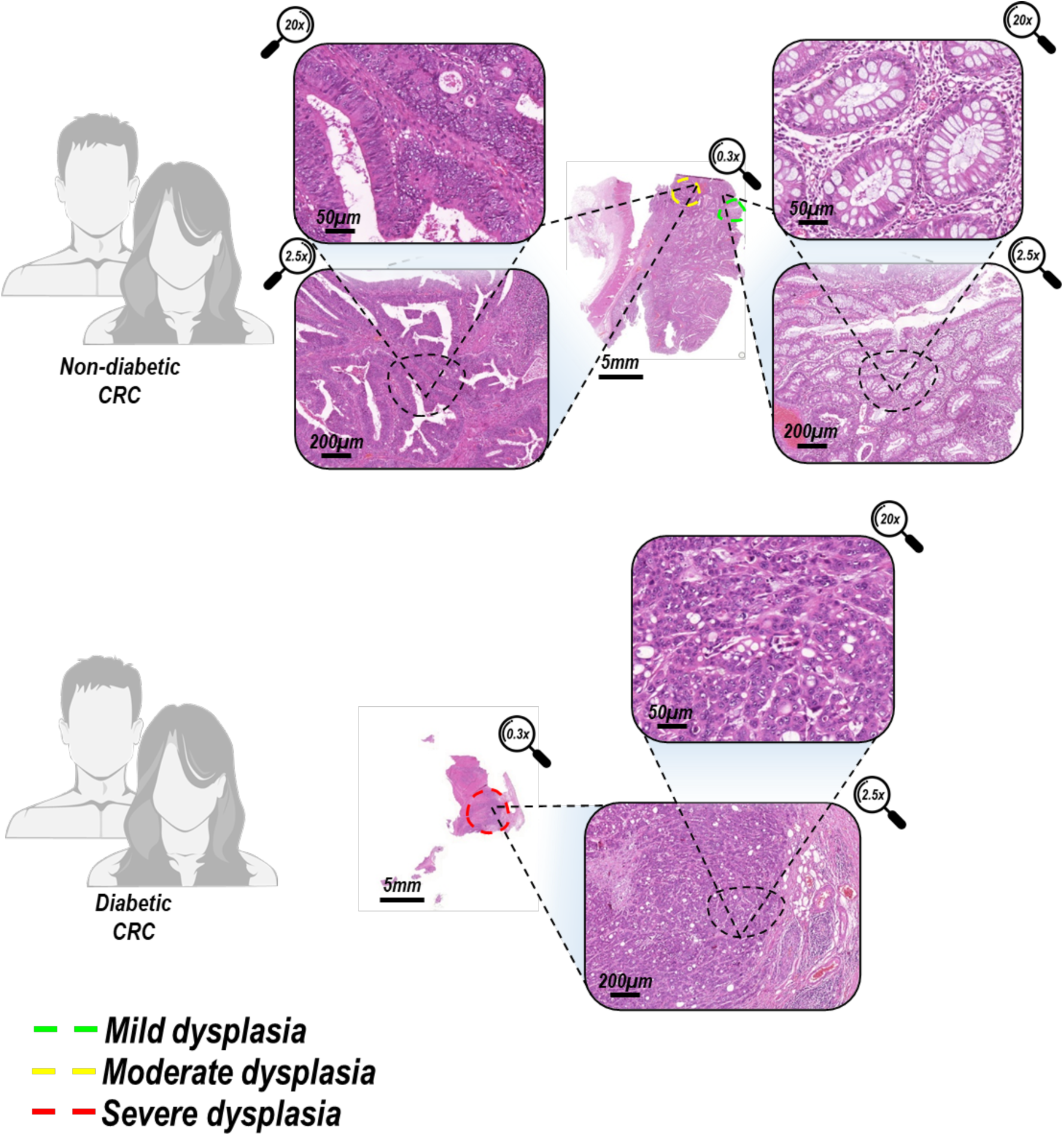
Histopathological alterations associated with CMAH deficiency and hyperglycemia in murine and human colorectal cancer. Representative H&E-stained colorectal tumor sections from non-diabetic and diabetic patients. Whole-slide images and representative higher-magnification views are shown. Dashed lines indicate the regions selected for magnification, whereas colored dashed outlines identify areas of mild (green), moderate (yellow), and severe (red) dysplasia. Scale bars: 5 mm, 200 μm, and 50 μm.

